# PIKfyve is required for phagosomal Rab7 acquisition and the delivery and fusion of early macropinosomes to phagosomes

**DOI:** 10.1101/2022.09.19.508510

**Authors:** James H. Vines, Catherine M. Buckley, Jason S. King

## Abstract

Phagosome maturation is tightly regulated to ensure efficient delivery of the complex arsenal of antimicrobial activities that kill and digest captured microbes. Like other endocytic pathways, phagosome maturation is regulated by a combination of Rab GTPases and phosphoinositide signalling lipids (PIPs) which define membrane identity and recruit specific effectors. PIKfyve is a PI-5 kinase, which converts PI(3)P to PI(3,5)P_2_ on endosomes. Disruption of PIKfyve results in severe defects in phagosomal maturation but the underlying mechanism remains unclear. Here, we use the model professional phagocyte, *Dictyostelium discoideum* to dissect the role of PIKfyve in the crucial first steps of phagosome maturation. We find that, although early Rab5 dynamics are unaffected, loss of PIKfyve prevents phagosomes from acquiring Rab7 by fusion with a pool of Rab7 and V-ATPase positive endosomes. By following PIP dynamics using our recently characterised PI(3,5)P_2_-probe SnxA, we delineate multiple subpopulations of Rab7-positive endosomes that fuse sequentially with phagosomes. We identify one of these as PI(3,5)P_2_-positive macropinosomes, which dock and fuse with phagosomes in a PIKfyve-dependent manner. We therefore show that *Dictyostelium* phagosomes primarily accumulate Rab7 by vesicular fusion rather than from a cytosolic pool, and that this requires PIKfyve. In particular PI(3,5)P_2_ defines a specific subset of fusogenic macropinosomes, which we propose enables content mixing and the efficient bulk delivery of lysosomal components to phagosomes.

## Introduction

The degradation of endocytic cargo is a highly regulated cellular process. Endosomes must acquire the correct markers with temporal accuracy to facilitate their maturation. This is achieved through sequential regulatory steps, which identify compartments in different stages of maturation. In this way, effector proteins and lysosomal compartments are recruited over the course of maturation to mediate the recycling or degradation of internalised material. Disruption of any one of these key regulators can result in aberrant endosomal trafficking and severe physiological defects.

Inositol phospholipids define specific compartment membranes during endosomal maturation. They contain a glycerol backbone, two non-polar fatty acid tails, and an inositol head group which can be phosphorylated at any of three positions (3, 4 or 5) to form one of eight different phosphoinositide (PIP) species. Each PIP can recruit different effector proteins, allowing for spatially segregated protein enrichment to different membranes. Interconversion of PIP species by the activity of a large family of phosphatases and kinases therefore allows specific effectors to be recruited at specific times throughout maturation (Bohdanowicz and Grinstein, 2013).

PIKfyve is a phosphoinositide 5-kinase that is recruited to early endosomes or the yeast vacuole via a FYVE (Fab1, YOTB, Vac1 and EEA1) domain, which specifically binds PI(3)P (Cabezas et al., 2006; Sbrissa et al., 1999; Yamamoto et al., 1995). There it phosphorylates PI(3)P to produce PI(3,5)P_2_, one of the least abundant and well understood PIPs. Across an array of organisms, disruption of PIKfyve causes a range of trafficking defects, including characteristic swollen endosomes and defects in lysosomal degradation (Buckley et al., 2019; Choy et al., 2018; de Lartigue et al., 2009; Dove et al., 2009; Ikonomov et al., 2001; Kim et al., 2014; Krishna et al., 2016; Nicot et al., 2006). Precisely how PIKfyve disruption leads to these phenotypes is unclear, although several PI(3,5)P_2_-activated ion channels on endosomal membranes have been identified, including TRPML1 and TPC2, which have been implicated in endosome-endosome fusion events (Dong et al., 2010; Leray et al., 2022; Samie et al., 2013; Wang et al., 2012).

The Rab family of small GTPases also play key roles in regulating endolysosomal traffic (Borchers et al., 2021). Early endosomes are marked by Rab5, which recruits effector proteins essential for early endosome fusion events (Henry et al., 2004; Lippuner et al., 2009), including Vps34, the class III PI-3 kinase responsible for PI(3)P synthesis. As maturation progresses, Rab5 is exchanged with the lysosomal marker, Rab7. This switch has been shown to be mediated by the evolutionarily conserved Mon1-Ccz1 complex, which both deactivates and dissociates Rab5, and recruits and activates Rab7 on endosomes (Kinchen and Ravichandran, 2010; Langemeyer et al., 2020; Nordmann et al., 2010). Accumulation of Rab7 on late endosomes promotes fusion with lysosomal compartments and its disruption results in severely perturbed lysosomal delivery to phagosomes (Rupper et al., 2001).

Our previous work showed that PIKfyve is critical for phagosome maturation in the soil-dwelling amoeba *Dictyostelium discoideum* (Buckley et al., 2019). *Dictyostelium* is a well-characterised professional phagocyte, which feeds through phagocytosis of bacteria or bulk uptake of media by the related process of macropinocytosis. These processes are well-conserved across evolution (Boulais et al., 2010; King and Kay, 2019), and the amenability of *Dictyostelium* to genetic manipulation, biochemical analysis and fluorescence microscopy studies make it a useful model system.

*Dictyostelium* mutants lacking PIKfyve (ΔPIKfyve) have severe defects in phagosomal proteolysis and killing (Buckley et al., 2019). These defects were, in part, caused by perturbed V-ATPase and hydrolase delivery to newly-formed phagosomes. Importantly, loss of PIKfyve renders *Dictyostelium* hypersensitive to infection with *Legionella pneumophila*, demonstrating the physiological importance of PIKfyve in innate immunity. PIKfyve is also important for phagosome-lysosome fusion in both macrophages and neutrophils (Dayam et al., 2015; Dayam et al., 2017; Isobe et al., 2019; Kim et al., 2014). This indicates a central and conserved role, but how PIKfyve activity is regulated and integrates with other components of the phagosomal maturation machinery such as Rab signalling and fusion with other endosomal compartments is poorly understood.

Here, we dissect how PIKfyve disruption causes defective phagosome maturation. By examining key maturation markers (Rab5, PI(3)P, V-ATPase, and Rab7) we find that although PIKfyve-deficient phagosomes acquire and recycle Rab5 normally, they accumulate almost no Rab7, which appears to be normally delivered by vesicle fusion. Using our recently described reliable reporter for PI(3,5)P_2_ (SnxA) (Vines et al., 2023), we identify an additional route for Rab7 enrichment via fusion with PI(3,5)P_2_-positive macropinosomes. We therefore show PIKfyve has a specific role in regulating heterotypic fusion between phagosomes and a temporally defined population of macropinosomes, which we suggest plays a previously unappreciated role in maintaining digestive efficiency.

## Materials and Methods

### Dictyostelium Culture

All *Dictyostelium discoideum* cells were derived from the Ax2 (Kay) laboratory strain background unless stated otherwise and grown in adherent culture in filter sterilised HL5 medium (Formedium) at 22^◦^C. Cells expressing extrachromosomal plasmids were transformed by electroporation and grown in appropriate antibiotic selection by addition of either 20 μg/mL hygromycin (Invitrogen) or 10 μg/mL G418 (Sigma). The PIKfyve knockout strain in Ax2 background was previously described (Buckley et al., 2019). Live cell imaging was performed in defined SIH medium (Formedium).

### Molecular biology

All gene sequences used for cloning were obtained from dictybase (www.dictybase.org) (Fey et al., 2013). The generation of SnxA-GFP (pJSK619), PIKfyve-GFP (pJV0025) and integrating SnxA-GFP plasmids has been described previously (Vines et al., 2023). GFP-Rab5A (pJV0054) was generated similarly: cDNA was cloned via PCR using primeSTAR max DNA polymerase (Clontech). This was followed by subcloning into a zero blunt TOPO II vector (Life Technologies), before cloning into the appropriate BglII/SpeI sites of the pDM expression plasmid (Paschke et al., 2018; Veltman et al., 2009). The coding sequence was confirmed by restriction digest. 2xFYVE-GFP (pJSK418) uses the sequence from the human Hrs gene and was previously described (Calvo-Garrido et al., 2014). GFP-Rab7A expression plasmid (pTH70) was kindly gifted by Huaquing Cai (Tu et al., 2022). Co-expression plasmids were created by first cloning the RFP-fusion sequence into the pDM series shuttle vectors (pDM1042 and pDM1121), before cloning as an NgoMIV fragment into the appropriate GFP-expression vector.

### Preparation of fluorescent yeast and dextran

*Saccharomyces cerevisiae* were used for phagocytosis assays. To obtain non-budded yeast, cells were grown for 3 days at 37^◦^C in standard YPD media (Formedium) until in stationary phase. For a budded population, growth was halted while yeast were still in log phase. Both populations were then centrifuged at 1000 x *g* for 5 minutes and resuspended to a final concentration of 1 × 10^9^ cells/mL in PBS pH7, then frozen until use. Yeast were fluorescently labelled using either pHrodo red succinimidyl ester (Life Technologies), or Alexa-405 succinimidyl ester (Life Technologies) at a final concentration of 0.25 mM and 2.5 mM, respectively. 0.5 × 10^9^ yeast were resuspended in 200 μL PBS at pH8.5 and incubated with 10 μL of prepared dye for 30-minutes at 37^◦^C with gentle shaking. After, yeast were pelleted and sequentially washed in 1 mL of PBS pH 8.5, 1 mL of 25nM TrisHCl pH8.5, and 1 mL of PBS pH 8.5 again. Finally, yeast were resuspended in 500 μL of KK2 pH 6.1 and kept at -20^◦^C. Yeast were diluted in SIH to a working concentration of 1 × 10^8^ yeast/mL before use. Texas-Red 70kDa Dextran (Life Technologies) was resuspended in water and diluted to a working concentration of 2 mg/mL. Far-red dextran was generated by labelling 70 kDa dextran (Invitrogen) with Alexa Fluor 680 NHS Ester (Invitrogen) as above, removing unbound dye by dialysis in KK2.

### Microscopy and image analysis

Approximately 1 × 10^6^ *Dictyostelium* cells were seeded into 35 mm microscopy dishes with glass bottoms (MatTek P35G-1.5-14-C) for fluorescence microscopy and left to grow overnight in SIH medium (Formedium). Imaging was conducted using a Zeiss LSM880 AiryScan Confocal microscope equipped with a 63× Plan Apochromat oil objective. To perform dual colour timelapse phagocytosis assays, most of the medium was removed just prior to imaging, and 20 μL of 1 × 10^8^ dyed yeast added. After 60-seconds, a thin layer of 1% agarose in SIH was overlaid on the cells, and excess medium was removed. For normal timelapses, images were captured every 10-seconds for up to 10-minutes. For extended timelapses, images were captured every 60-seconds. Fluorescent intensity analysis around each phagosomal membrane was carried out automatically using the Python plugin pyimagej, which has been described previously (Vines et al., 2023). In brief, phagosomal contents were identified and segmented by using the fluorescent yeast, which are large and easy to follow. The fluorescence channel containing the yeast was thresholded, and large yeast-sized particles identified to be followed over time by examining similar particles on adjacent frames. This allowed individual phagosomes to be followed over the course of a few minutes. Increases in membrane fluorescence were then calculated by expanding the particle perimeter and taking average fluorescence readings in a banded area.

To perform dextran pulse-chases, the majority of the medium was removed and replaced with 50 μL of 2 mg/mL 70kDa Texas Red dextran for the specified duration. Dextran was then removed by washing three times in SIH medium, leaving ∼1 mL of SIH in the dish. Cells were immediately imaged, with multiple fields of view captured every 2-minutes for 10-minutes. Quantification of macropinosome number and size was quantified using an automated imageJ script. GFP co-localization at each time point was scored manually. For macropinosomes-phagosome fusion assays, 50 μL of 2 mg/mL Texas Red dextran was added to cells for the time indicated for macropinosomes to form. The dish was then washed as above, before addition of 20 μL of 1 × 10^8^ dyed yeast and then agarose overlay after a further 60-seconds. Clustering analysis was performed using a modified version of the automated analysis pipeline described above. For macropinosome/macropinosome mixing a second dye, 50 μL of Alexa680 70kDa Dextran, was added instead of yeast. For macropinosome/phagosome fusion assays, budded yeast were used instead and analysis of fusion performed manually in ImageJ by measuring the mean intensity across the yeast neck over time. For analysis of clustering macropinosomes, macropinosomes within 1 μm of the phagosome were manually scored as GFP positive or negative, and binned into 2-minute intervals.

### Western Blotting

Wild-type and ΔPIKfyve cells expressing GFP-Rab5A, GFP-2xFYVE, or GFP-Rab7A were analysed by SDS-PAGE and Western blot using a rabbit anti-GFP primary antibody (gift from Andrew Peden) and a fluorescently conjugated anti-rabbit 800 secondary antibody, using standard techniques. The endogenous biotinylated mitochondrial protein methylcrotonoyl-CoA Carboxylase 1 was used as a loading control using Alexa680-conjugated streptavadin (Davidson et al., 2013).

### Statistics

Graphpad Prism 9 was used for statistical analysis. The figure legends provide details on the number of biological replicates and the statistical tests used for each experiment. A p-value of <0.05 was considered significant, with * indicating p>0.05, ** indicating p>0.01, and *** indicating p>0.005 throughout.

## Results

### PIKfyve is required for Rab7 delivery to phagosomes

To understand the functional role of PIKfyve, we examined how its loss affected other core components of the phagosome maturation pathway. Previously, we observed that PIKfyve-GFP is transiently recruited to *Dictyostelium* phagosomes between 30 to 120-seconds following engulfment (Vines et al., 2023), so we focused our analysis on components likely to be active within this time frame. During classical endocytosis, Rab5 accumulates at early stages and exchanges with Rab7 as endosomes mature (Rink et al., 2005). A similar Rab5-Rab7 exchange has been described during both phagosome and macropinosome maturation around the same time we observe PIKfyve-GFP recruitment (Kerr et al., 2006; Poteryaev et al., 2010; Tu et al., 2022; Vieira et al., 2003).

Co-expression of RFP-Rab5A and GFP-Rab7A allowed us to follow the dynamics of both proteins to newly formed phagosomes in wild-type and ΔPIKfyve cells (**Figure 1A-B** and **Video 1**). In wild-type cells, as previously reported (Tu et al., 2022), RFP-Rab5A was highly enriched on perinuclear endosomes and to a lesser extent at the plasma membrane. RFP-Rab5A was further recruited to phagosomes following internalisation before its removal in the following minutes. As expected, this coincided with a gradual accumulation of GFP-Rab7A, which also localized to a large pool of small endosomes which continuously clustered around phagosomes and appeared to fuse from approximately 30 seconds post-engulfment. In contrast, although RFP-Rab5A dynamics appeared identical in ΔPIKfyve cells, phagosomes only ever accumulated very low levels of GFP-Rab7A, despite strong recruitment to other enlarged endocytic compartments (**Figure 1B**).

**Figure 1:**
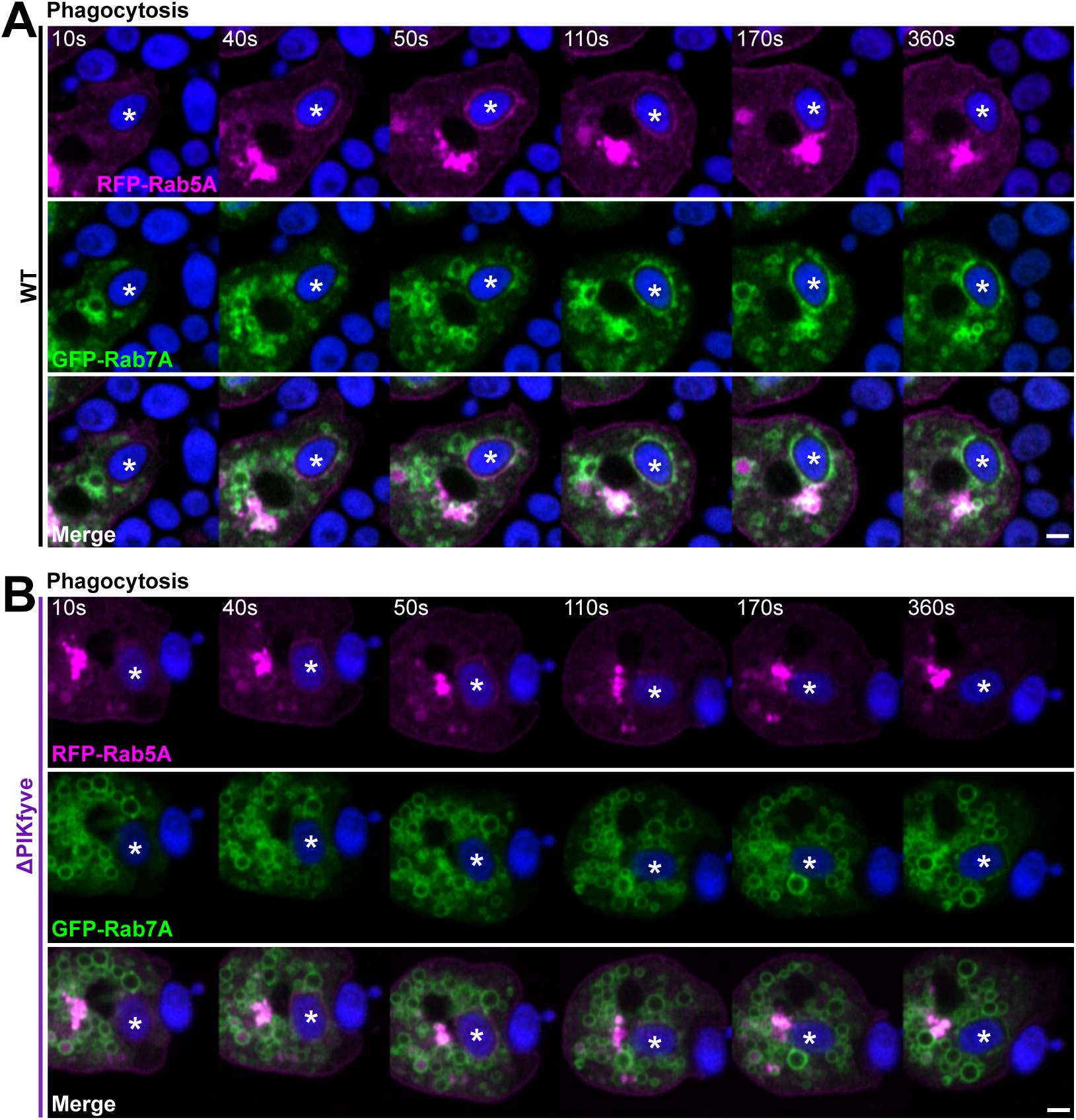
Simultaneous analysis of RFP-Rab5 and GFP-Rab7 recruitment. Timelapse movies of (A) Wild-type and (B) ΔPIKfyve cells co-expressing RFP-Rab5A (magenta) and GFP-Rab7A (green) during phagocytosis of Alexa405-yeast (blue). Rab7A only arrives on wild-type phagosomes. The followed phagosome is indicted by an asterisk. The full timeseries is shown in Video 1. All scale bars = 2 μm

To quantify changes in protein enrichment around engulfed particles over multiple events, we employed an automated image analysis pipeline which we have previously described (Vines et al., 2023). This segments the yeast and measures the intensity of fluorescent fusion proteins around the periphery at each time point. To reduce the possibility of artefacts from overexpressing both proteins during quantification, Rab5A and Rab7A were expressed individually as GFP-fusions. Multiple events were then averaged and normalised to the intensity at the moment engulfment completed (**Figure 2**).

**Figure 2:**
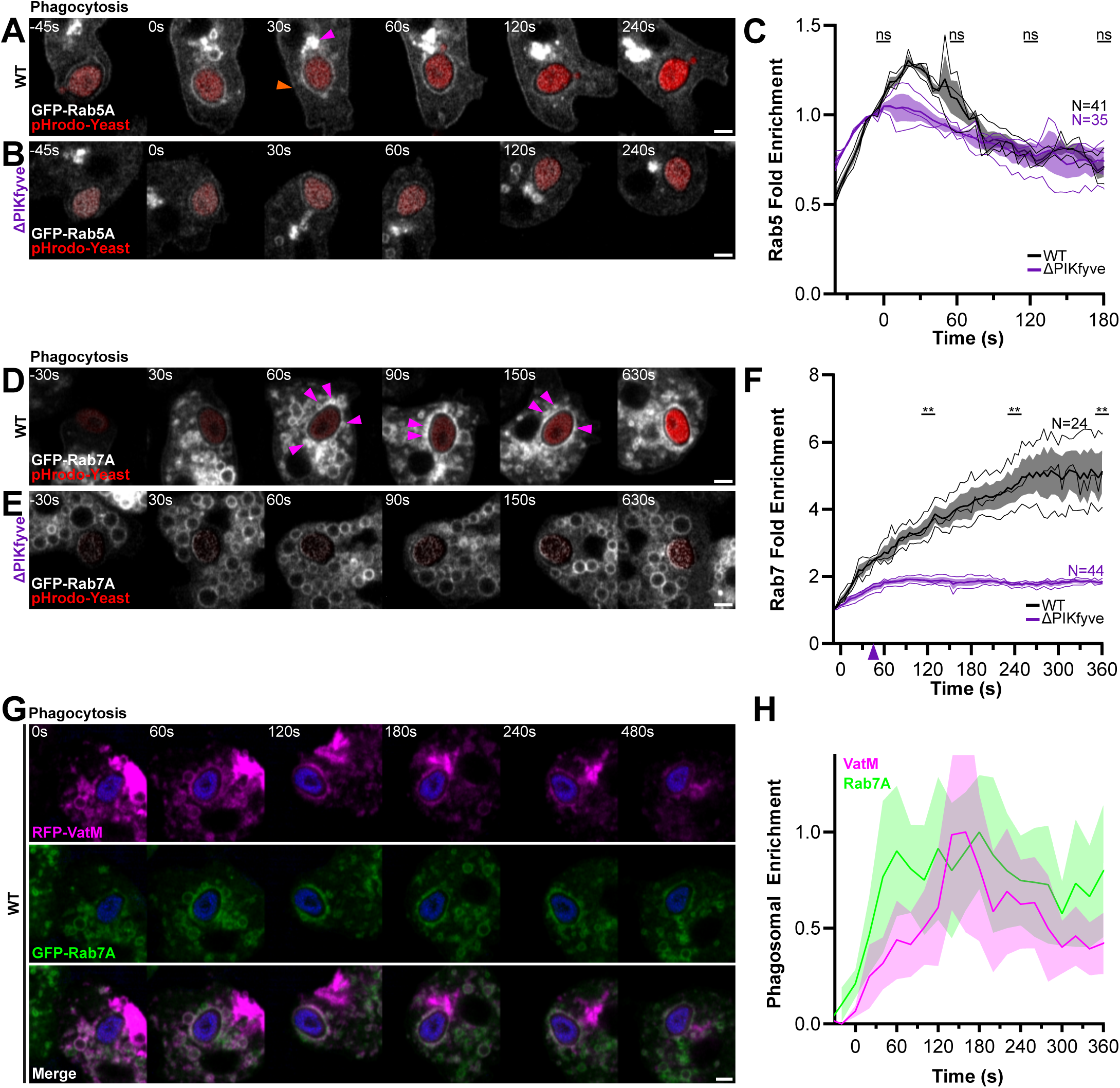
Loss of PIKfyve affects Rab7 recruitment to phagosomes. (A) and (B) Timelapses of GFP-Rab5A during phagocytosis of pHrodo-yeast (red) in wild-type (A) and ΔPIKfyve (B) cells. GFP-Rab5A is constitutively enriched on a perinuclear pool (pink arrow) and the plasma membrane (orange arrow) as well as the phagosomal envelope for 0s-60 s. See Video 2 for complete sequence. Quantification of GFP-Rab5A enrichment averaged over the indicated number of independent events is shown in (C). No significant differences were found at any timepoint tested (paired T-test). (D) and (E) show GFP-Rab7A recruitment to phagosomes in wild-type and ΔPIKfyve cells respectively. Arrowheads indicate the clustered GFP-Rab7A-enriched vesicles only observed in wild-type cells. See Video 5 for complete sequence. (F) Quantification of phagosomal GFP-Rab7A enrichment, demonstrating a significant decrease in ΔPIKfyve mutants. (G) Timelapse movie of wild-type cells co-expressing RFP-VatM (magenta) and GFP-Rab7A (green). Fluorescence intensity around phagosome in G is shown in (H). Data is mean intensity ± SD of a linescan along the phagosomal membrane normalised between minimum and maximum intensities observed. All scale bars = 2 μm.

This analysis confirmed the enrichment of GFP-Rab5A starting just prior to phagocytic cup closure (**Figure 2A-C** and **Video 2**). This was independent of successful internalisation (**Video 3**), suggesting a potential role for Rab5 before engulfment is complete in *Dictyostelium*. Rab5A-GFP then intensified following internalisation and peaked 30-seconds after engulfment (**Figure 2A**). Although expression levels were lower in the mutants (**Figure S1A**), the timing of Rab5A-GFP recruitment was not significantly changed in ΔPIKfyve cells (**Figure 2B-C**).

As Rab5 is also responsible for PI(3)P synthesis, due to the recruitment and activation of the Class 2 PI-3 kinase Vps34, we also quantified PI(3)P using the well characterised reporter GFP-2xFYVE (Gillooly et al., 2000). In wild-type cells, GFP-2xFYVE became enriched on phagosomes in a uniform ring within 60 seconds of engulfment (**Figure S1B-D** and **Video 4**) and was maintained for at least 50 minutes (**Figure S1E**). PI(3)P also localized to a set of small vesicles throughout the cytosol (arrows in **Figure S1B**). In ΔPIKfyve cells, however, these vesicles were replaced with large numbers of characteristically swollen endosomes which moved slowly around the cell (**Figure S1C**). Nonetheless, there were no quantifiable differences in phagosomal PI(3)P dynamics in ΔPIKfyve cells, consistent with normal Rab5 dynamics (**Figure S1D**).

Quantification of GFP-Rab7A in wild-type cells showed a continuous accumulation to phagosomes starting around 30 seconds post-engulfment and continuing for the next ∼5 minutes (**Figure 2D-F** and **Video 5**). GFP-Rab7A was then maintained for at least 50 minutes (**Figure S1F-G**), most likely until the transition to a post-lysosome (Buczynski et al., 1997). In ΔPIKfyve cells however, quantitative analysis confirmed the majority of phagosomal GFP-Rab7A was absent, although very low levels of GFP-Rab7A were visible at some timepoints (**Figure 2F**).

In these experiments, it was notable that while GFP-2xFYVE appeared as a smooth ring directly on the phagosomal membrane, in wild-type cells GFP-Rab7A-positive vesicles appeared to cluster around the phagosome before fusing. This was apparent by analysis of the variance in fluorescence along the phagosome membrane (**Figure S1H**) and indicates that phagosomal Rab7 accumulation is predominately driven by fusion with other Rab7-containing compartments, rather than recruitment from a cytosolic pool. In contrast, although the cytosol of ΔPIKfyve cells was full of swollen GFP-Rab7A positive vesicles, there was no obvious clustering around the nascent phagosome (**Figure 2D**).

We have previously shown that disruption of PIKfyve causes a reduction in both V-ATPass and lysosomal enzyme delivery to phagosomes (Buckley et al., 2019). We therefore asked whether Rab7 and V-ATPase were delivered on the same vesicles by co-expressing GFP-Rab7A with RFP-VatM (a component of the V-ATPase complex) in wild-type cells (**Figure 2G**). This showed that all VatM-positive vesicles were also Rab7A-positive, and both proteins accumulated on the phagosomal membrane concurrently (**Figure 2H**). This indicates that PIKfyve is required for the delivery of Rab7A/VatM-positive endosomes to the early phagosome and its absence causes a generalized loss in the delivery of the lysosomal components required for maturation.

### PIKfyve is not important for Rab7 delivery to macropinosomes

Although GFP-Rab7A did not accumulate on ΔPIKfyve phagosomes, it clearly still localised to other large endocytic compartments (**Figure 2E**). As these resemble the swollen macropinosomes previously observed in these cells (Michael et al., 2013), we asked whether the defect in GFP-Rab7A delivery was specific to phagosome maturation. As individual macropinosomes are difficult to follow for more than 2-3 minutes by timelapse microscopy, we labelled macropinosomes with a 2-minute pulse of TexasRed-dextran before washing out into normal medium (**Figure 3A-B**). Different fields of view were then captured at time intervals after dextran removal allowing the percentage of macropinosomes positive for GFP to be measured for up to 10 minutes (Buckley et al., 2016). In this assay, GFP-Rab7A recruitment to macropinosomes was indistinguishable between ΔPIKfyve cells and wild-type controls, accumulating within 2 minutes of internalisation (**Figure 3C**).

**Figure 3:**
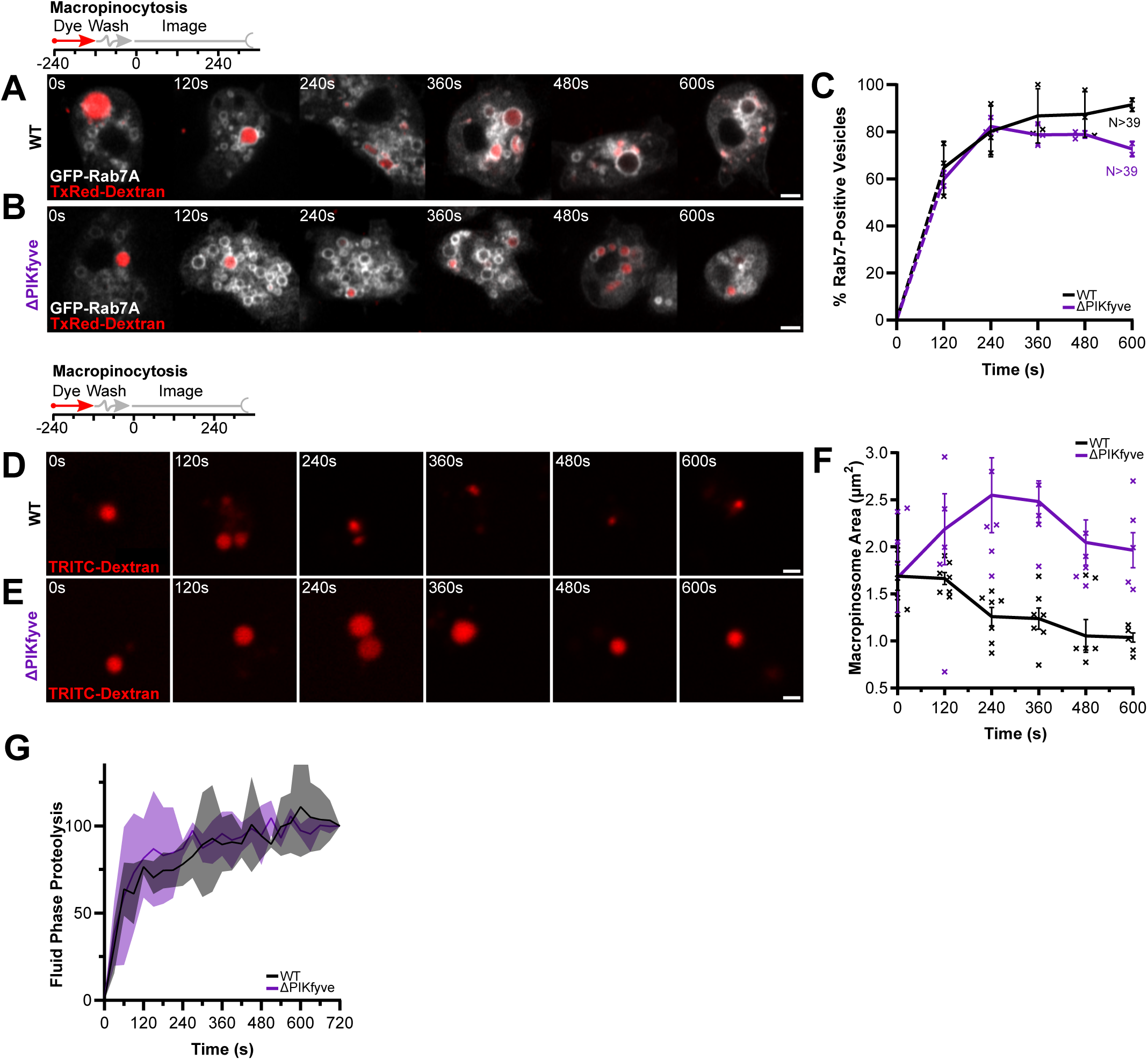
PIKfyve is not important for Rab7 delivery to macropinosomes. (A) and (B) GFP-Rab7A recruitment to macropinosomes labelled by a 2-minute pulse of TxRed dextran (red) in wild-type (A) and ΔPIKfyve(B) cells. (C) Quantification by scoring the percentage of GFP-positive macropinosomes at each time point after dextran wash out. GFP-Rab7A dynamics are similar in both cell types. Data shown is mean ± SEM of 3 independent experiments, with N indicating the total number of cells measured. Biological replicates shown by crosses. (D) and (E) labelling of macropinosomes by a 2-minute pulse of TRITC dextran (red) in wild-type and ΔPIKfyve cells respectively. (F) Quantification of macropinosome shrinkage during early maturation. Wild-type macropinosomes lose approximately half their volume within 5 minutes, but this is disrupted in ΔPIKfyve cells. Graph represents the mean ± SEM of six independent experiments. Biological replicates shown by crosses. (G) Fluid phase proteolytic activity in wild-type and ΔPIKfyve cells measured by increase in fluorescence after a 2-minute pulse of DQ-BSA. Data shown is mean ± SD of three independent experiments. All scale bars = 2 μm.

Although GFP-Rab7A dynamics appeared normal, the macropinosomes in ΔPIKfyve cells still appeared much larger. Quantification of their size and fluorescence intensity demonstrated that although macropinosomes started off the same size, the normal shrinkage and concentration of their lumen was significantly impaired in PIKfyve mutants (**Figure 3D-F**). Therefore shrinkage and Rab7 accumulation are functionally separable, indicating multiple independent roles of PIKfyve.

To functionally assess macropinosome maturation, we also measured proteolysis of the reporter dye DQ-BSA over time. Although loss of PIKfyve almost completely blocks proteolysis of DQ-BSA when conjugated to beads (Buckley et al., 2019), degradation of the fluid-phase dye was completely unaffected (**Figure 3G**). Therefore, despite generally similar maturation pathways, PIKfyve plays a specific role in the fusion of lysosomes to phagosomes, but not macropinosomes.

### PI(3,5)P_2_ is delivered by fusion of small Rab7-positive vesicles

The evidence above suggests that PIKfyve or production of PI(3,5)P_2_ is required for lysosomal fusion with phagosomes. To better understand how Rab7 is recruited relative to PI(3,5)P_2_, we co-expressed RFP-Rab7A with PIKfyve-GFP in ΔPIKfyve cells (**Figure 4A**). As expected, PIKfyve-GFP expression restored Rab7A recruitment to phagosomes and recruitment of PIKfyve-GFP overlapped with the start of Rab7 enrichment (**Figure 4B**). As we recently showed, PIKfyve recruitment was only transient and was lost from the phagosomal membrane within 2 minutes (Vines et al., 2023). The sustained presence of its product PI(3,5)P_2_ is therefore likely to drive continued Rab7 recruitment.

**Figure 4:**
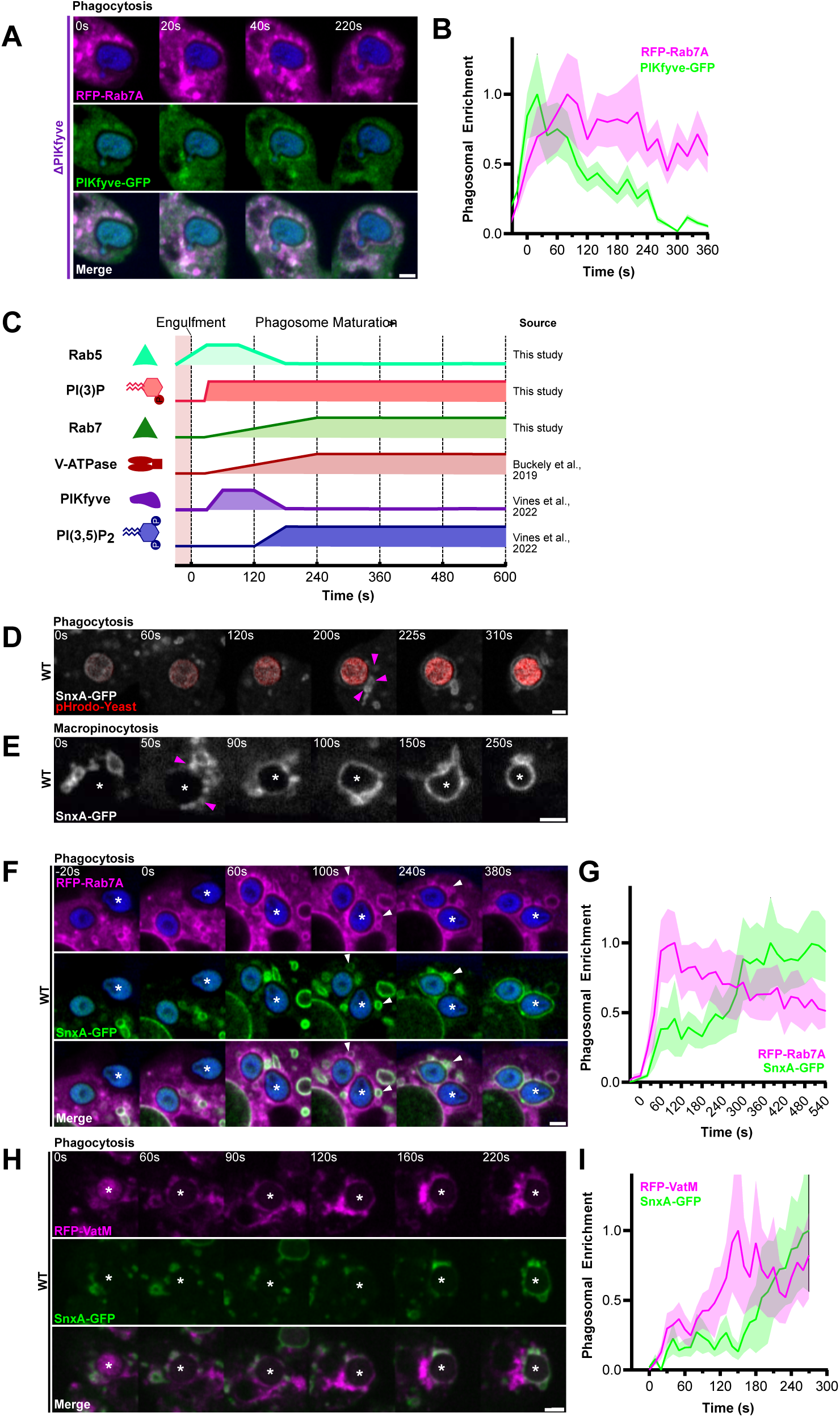
PI(3,5)P_2_ is delivered after Rab7 to both macropinosomes and phagosomes. (A) Timelapse of RFP-Rab7A and PIKfyve-GFP in ΔPIKfyve cells (rescued). Fluorescence intensity around phagosome in (A) is shown in (B). (C) Summary of phagosome maturation dynamics determined from this, and previous studies. When the line is low, the respective component is absent from the phagosomal membrane, when the line is high the component is enriched. (D) and (E) Timelapses of SnxA-GFP recruitment following either phagocytosis of a pHrodo-yeast (red) or macropinosomes formation respectively. Pink arrowheads indicate clustering SnxA vesicles. See Video 6 for full sequence. (F) Timelapse of RFP-Rab7A (magenta) and SnxA-GFP (green) during phagocytosis of Alexa405-yeast (blue) in wild-type cells. Arrowheads indicate clustered vesicles marked by both SnxA and Rab7. Fluorescence intensity around this phagosome is shown in (G). (H) Timelapses of RFP-VatM (magenta) and SnxA-GFP (green) during phagocytosis in wild-type cells. Fluorescence intensity around phagosome in (H) is shown in (I). For all intensity graphs, data is mean intensity ± SD of a linescan along the respective phagosomal membrane normalised between minimum and maximum intensities observed. All scale bars = 2 μm.

To observe PI(3,5)P_2_ dynamics, we used our recently identified biosensor, SnxA, which specifically binds this lipid with high affinity (Vines et al., 2023). In *Dictyostelium*, we showed SnxA-GFP is enriched on both phagosomes and macropinosomes approximately 120-seconds after engulfment, just before PIKfyve itself leaves the phagosome. Upon disruption of PIKfyve, SnxA-GFP becomes completely cytosolic. For clarity, the dynamics of PI(3,5)P_2_ during phagosome maturation relative to other markers from this and other studies is summarised in **Figure 4C**.

Closer examination of SnxA-GFP recruitment to phagosomes and macropinosomes by timelapse microscopy, indicated that accumulation of PI(3,5)P_2_ is accompanied by the docking and fusion of multiple smaller SnxA-GFP-positive vesicles less than 1 μm in size. These cluster around macropinosomes and phagosomes several seconds before they acquire SnxA-GFP themselves (arrows in **Figure 4D-E** and **Video 6**). This implies that PI(3,5)P_2_ accumulates through both *in situ* synthesis by PIKfyve and delivery via fusion with another PI(3,5)P_2_ compartment. This suggests a role for PIKfyve in regulating the docking or fusion of a specific subset of vesicles to the phagosome.

We then asked whether the small PI(3,5)P_2_-positive vesicles also contained Rab7, by co-expressing SnxA-GFP with RFP-Rab7A. This demonstrated that all docked SnxA-positive vesicles were also positive for Rab7A (White arrows **Figure 4F** and **Figure S2A**). However, RFP-Rab7A also localised to additional populations of vesicles that start to cluster and fuse much earlier than those marked by SnxA, as is evident in the quantification of intensity (**Figure 4G**). Co-expression of SnxA-GFP alongside RFP-VatM yielded similar results, with SnxA vesicles representing a later-fusing subset of VatM-positive endosomes (**Figure 4H-I**).

Together, our data demonstrate the presence of a Rab7/VatM/PI(3,5)P_2_-positive compartment which is delivered to early phagosomes following PIKfyve recruitment, approximately 120 seconds post-engulfment. This compartment is separate from an additional population of Rab7/V-ATPase-positive endosomes which cluster earlier and do not contain PI(3,5)P_2_. This indicates the sequential delivery of lysosomal components and depends on PIKfyve for its regulation.

### The clustering PI(3,5)P_2_-positive vesicles are macropinosomes

We next investigated the origin of the pool of Rab7/V-ATPase/PI(3,5)P_2_-positive vesicles that fuse with phagosomes. Our previous work shows the primary compartment marked by SnxA in *Dictyostelium* and mammalian cells are macropinosomes (Vines et al., 2023) (**Figure 4E**) and this temporally overlaps with the accumulation of Rab7 we show here (**Figure 3A**). Others have also shown that macropinosomes can undergo homotypic fusion; (Dolat and Spiliotis, 2016; Hamasaki et al., 2004; Neuhaus et al., 2002; Tu et al., 2022) using sequential pulses of different coloured dextrans, we were also able to observe macropinosomes fusing with one another from relatively early stages of maturation (**Figure S2B**). Therefore, we hypothesised that Rab7/V-ATPase/PI(3,5)P_2_ macropinosomes fuse with phagosomes in a PIKfyve-dependent manner.

To test this, we labelled macropinosomes by incubating cells with a 5 minute pulse of 70 kDa dextran, before washing and addition of yeast. In this way, we were able to follow interactions between a defined population of dextran-filled macropinosomes and nascent phagosomes. Using cells co-expressing 2xFYVE-RFP and SnxA-GFP, we observed that the SnxA-positive vesicles clustering around nascent phagosomes also contained dextran and were positive for RFP-2xFYVE (white arrows **Figure 5A**). We previously showed that both lipids are only present on macropinosomes between 3-8 minutes of maturation (King and Kay, 2019; Vines et al., 2023). This matches the timings of our pulse-chase experiment and indicates macropinosomes cluster phagosomes at a defined point in their maturation.

**Figure 5:**
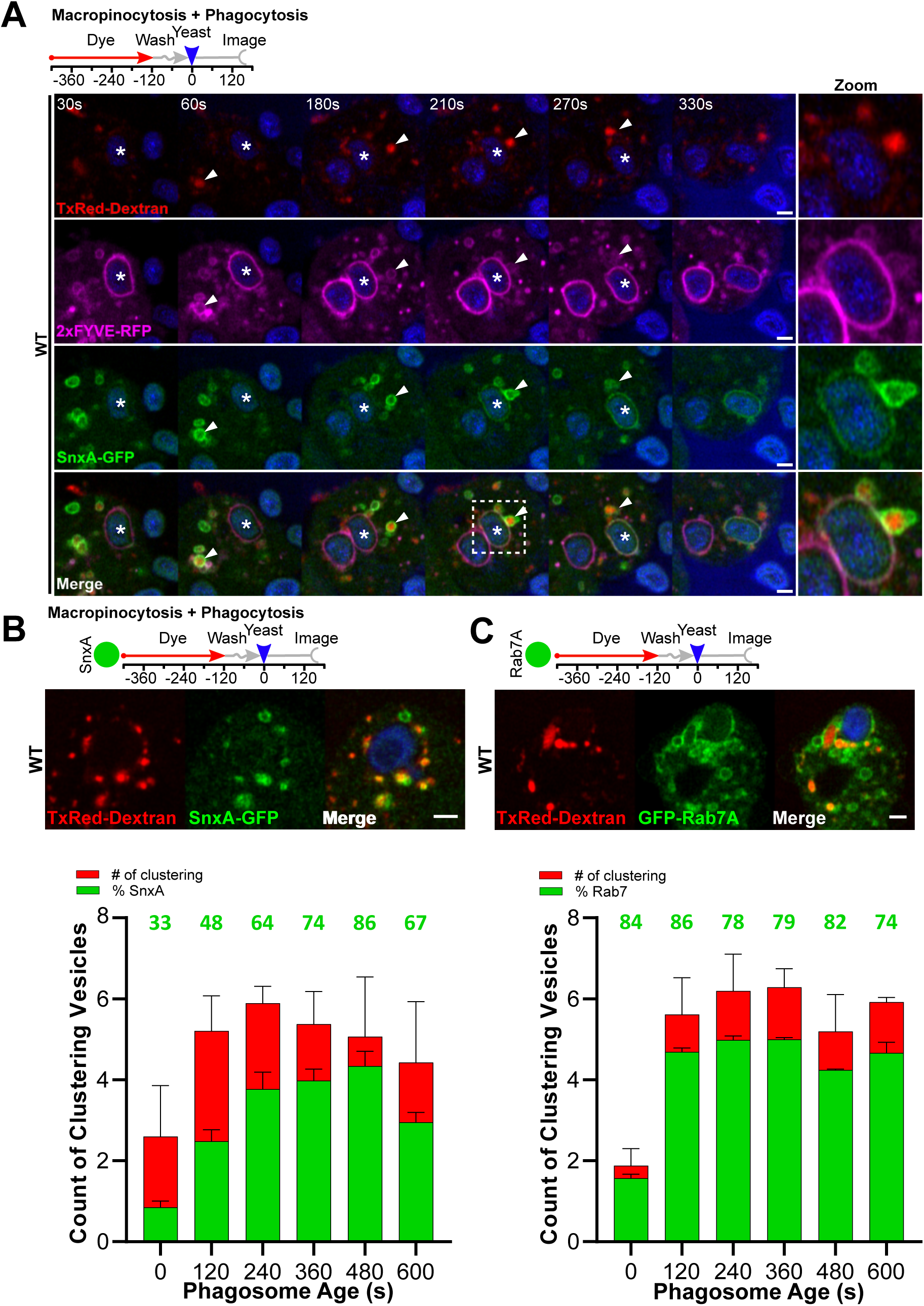
Rab7/PI(3)P/PI(3,5)P_2_ positive compartments cluster around macropinosomes. (A) Timelapse of cells co-expressing 2xFYVE-RFP (magenta) and SnxA-GFP (green) during phagocytosis of Alexa405-yeast (blue) in wild-type cells. Schematic shows experimental procedure: Cells were pre-incubated for 5 minutes with a 70kDa Txred-dextran (red) to load macropinosomes before dextran washout and addition of yeast. White arrow indicates a 2xFYVE and SnxA positive macropinosome which docks and fuses with the phagosome during acquisition of SnxA to the phagosomal membrane. (B and C) Images and quantification of clustering macropinosome identity. Wild-type cells expressing GFP-Rab7A or SnxA-GFP were treated with a dextran pulse-chase before phagocytosis of yeast. Macropinosomes within 1 μm of the phagosome were identified at each timepoint and both total number (red bars) and proportion colocalizing with the respective GFP-reporter (green bars and numbers) were scored manually. Frames were binned into 2-minute groups, data shown are the mean ± SEM of three independent experiments. All scale bars = 2 μm.

In addition to SnxA-positive macropinosomes, we also observed additional populations of SnxA-GFP negative macropinosomes, which clustered earlier. To determine the identities of these macropinosomes and how they changed over time, we imaged macropinosome clustering in cells expressing different GFP-fusion reporters as above. At 2 minute intervals post-phagocytosis, all dextran-labelled macropinosomes within 1 μm of the phagosomal membrane were scored for the presence of each reporter (**Figure 5B-C**). In each experiment, the number of phagosome-proximal macropinosomes started low (initially ∼2 per phagosome) but increased to ∼6 per phagosome by 2 minutes, indicating temporal regulation of the interaction between the two compartments.

Consistent with our timelapses of SnxA-GFP delivery, only 33% of proximal macropinosomes were positive for SnxA-GFP at initial timepoints. Based on the timelapse experiments above, this likely indicates the background false-positive rate for this method caused by macropinosomes that are near phagosomes by chance. This increased to 80% at 6 and 8 minutes, consistent with the SnxA-GFP-positive macropinosome docking observed above. In contrast, at all timepoints 80% of proximal macropinosomes were positive for GFP-Rab7A. Therefore the increased population of macropinosomes that actively cluster around phagosomes around 5 minutes is defined by PI(3,5)P_2_, rather than Rab7.

### PIKfyve is required for fusion of macropinosomes with phagosomes

As the clustering of macropinosomes correlates with the presence of PI(3,5)P_2_, we tested whether this was perturbed by PIKfyve disruption. For this, we repeated the dextran and yeast pulse-chase described above in both wild-type and ΔPIKfyve cells. As before, in wild-type cells, macropinosomes began clustering around phagosomes from within 30 seconds of engulfment (**Figure 6A**). This was completely lost in ΔPIKfyve cells, where the enlarged macropinosomes were never clearly observed interacting with the phagosome (**Figure 6B**). We quantified this by measuring the average dextran signal within 0.5 μm of the yeast, which indicated that this failure to cluster was persistent throughout maturation (**Figure 6C**).

**Figure 6:**
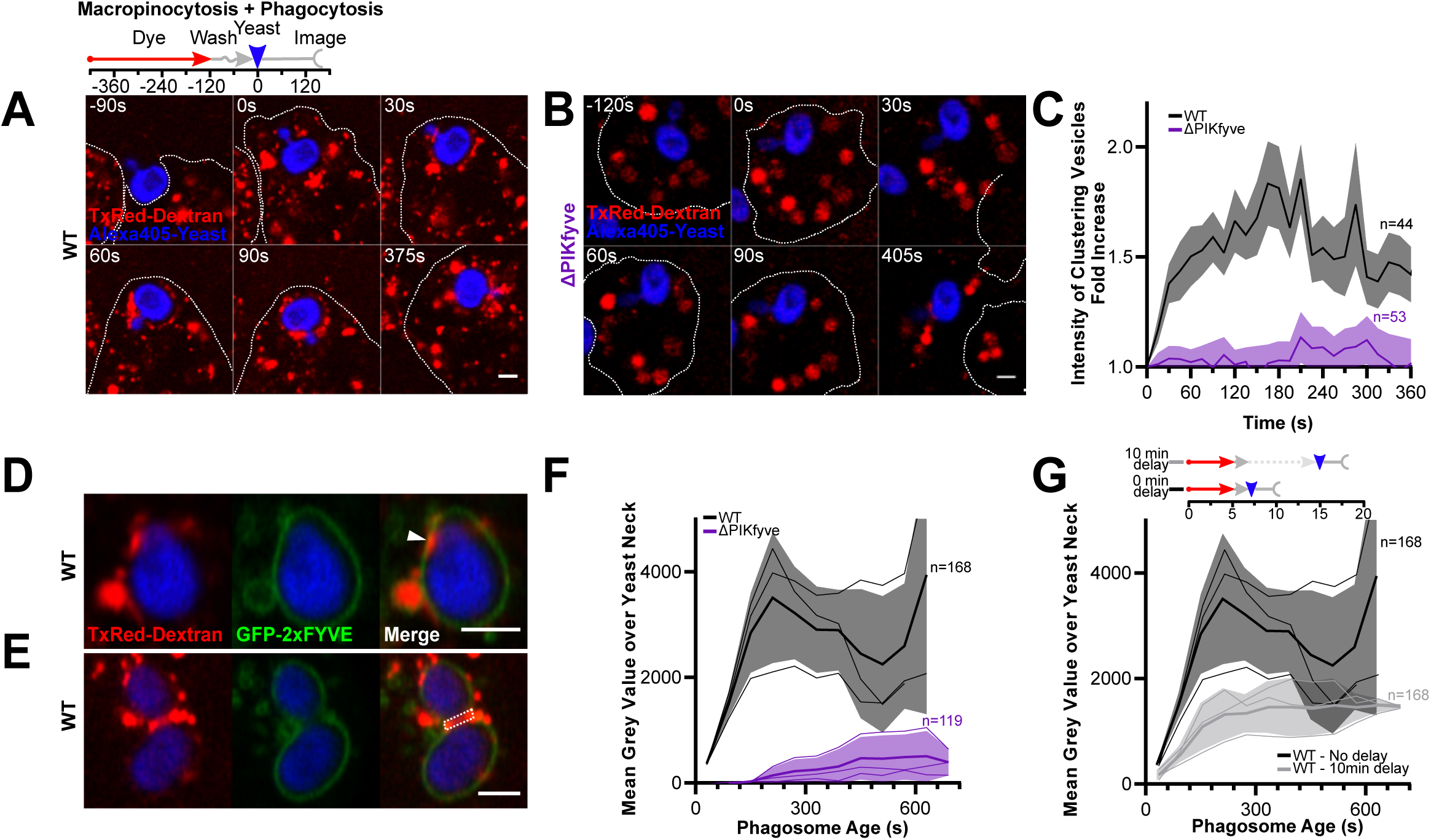
PIKfyve is required for macropinosome-phagosome fusion. (A) and (B) Timelapses of interactions between early macropinosomes (red) and phagosomes (blue) in wild-type and ΔPIKfyve cells, after sequential addition of fluorescent dextran and yeast. Schematic shows experimental procedure. Macropinosomes cluster around phagosomes in wild-type but not in ΔPIKfyve cells. (C) Quantification of dextran intensity within 0.5 μm of the phagosome, averaged over multiple events. Graph shows fold enrichment normalised to 10s before engulfment ± SEM. (D) TxRed-dextran (red) accumulation within the phagosomal membrane (green) marked by 2xFYVE (arrowhead). (E) Shows the same, using a budded yeast. Note strong dextran accumulation around the bud neck (dashed box). (F) Quantification of dextran intensity in this area averaged across multiple events ± SEM, binned by the age of the phagosome. (G) Quantification of dextran delivery (as in F) after a 10-minute delay before addition of yeast (see schematic, same control data as in F). Older macropinosomes show less delivery to phagosomes. All scale bars = 2 μm.

To determine whether macropinosomes were just clustering around phagosomes or fusing with them, we also examined delivery of dextran to the phagosomal lumen. Using GFP-2xFYVE to highlight the bounding phagosomal membrane, it was occasionally possible to observe dextran accumulation in small pockets that appeared to be within the phagosomal envelope (arrow, **Figure 6D**). This was difficult to reliably quantify, but we noticed that when budding yeast were engulfed, dextran preferentially accumulated within the bud neck region of the phagosome due to its negative curvature (**Figure 6E**). Using budded yeast therefore enabled us to reproducibly observe and quantify dextran delivery within the phagosome. This demonstrated fusion of early macropinosomes to phagosomes in wild-type cells, which was completely lost in ΔPIKfyve cells (**Figure 6F**). Therefore, PIKfyve is required for both clustering and delivery of macropinosomes to early phagosomes.

As both PI(3)P and PI(3,5)P_2_ are only present on macropinosomes for the first 7-8 minutes of maturation (Vines et al., 2023), we asked if macropinosomes were only fusion-competent over this period. To test this, cells were again exposed to a 5 minute pulse of dextran, but were then incubated for 10 minutes for the labelled macropinosomes to mature to a later stage before addition and phagocytosis of yeast. This delay significantly decreased accumulation of dextran at the yeast neck with only ∼20% of the dextran delivery remaining (**Figure 6G**). As macropinosomes also fuse with each other (**Figure S2B**), this residual macropinosome-phagosome fusion can potentially be attributed to older macropinosomes fusing to newer macropinosomes which are then able to fuse with the nascent phagosome. Macropinosomes are therefore only competent to fuse with phagosomes at a specific early stage (2-8 minutes old), when they possess both PI(3)P and PI(3,5)P_2_.

Our data demonstrate a specific role for PIKfyve in the delivery of macropinosomes to newly formed phagosomes. These macropinosomes are enriched in V-ATPase, Rab7 and lysosomal enzymes, all of which fail to accumulate on or in phagosomes in ΔPIKfyve cells (Buckley et al., 2019). We therefore propose fusion with macropinosomes provides a mechanism for the rapid delivery of lysosomal components to early phagosomes, which is physiologically important for effective bacterial killing and protection from pathogens (Buckley et al., 2016).

## Discussion

In this study we expand our understanding of the complex dynamics of Rab and phosphoinositide signalling that underpin the processing of phagosomes and macropinosomes. We describe a complex pathway in which distinct pools of endosomes are sequentially delivered to maturing phagosomes over the first few minutes of maturation. We find a portion of these endosomes are early macropinosomes which cluster around phagosomes before heterotopic fusion. While fusion of phagosomes with lysosomes is well studied (Dayam et al., 2015; Dayam et al., 2017; Isobe et al., 2019; Kim et al., 2014), fusion with macropinosomes is much less well understood. However, studies in both *Dictyostelium* and mammalian cells have also shown early macropinosomes fuse with one another in the first few minutes after formation, although it is unclear whether aged macropinosomes also possess this ability (Hamasaki et al., 2004; Schink et al., 2021).

Fusion between macropinosomes and phagosomes has also been observed in mammalian cells and is implicated during infection. Both *Salmonella enterica* and *Shigella flexineri* induce macropinosome formation from the ruffles formed during their entry into non-phagocytic host cells (Adam et al., 1996; Francis et al., 1993). These macropinosomes cluster around the bacteria-containing vacuole and appear to facilitate the rupture and escape of *Shigella* (Weiner et al., 2016), whereas they appear to stabilise and support the vacuolar lifestyle of *Salmonella* (Stevenin et al., 2019). Interestingly, inhibition of PIKfyve also disrupts Salmonella replication (Kerr et al., 2010). Although it is unclear how infection-associated macropinosomes relate to those made constitutively by professional phagocytes like macrophages or *Dictyostelium*, this highlights a common role for macropinosome delivery in the remodelling of microbe-containing compartments.

The observation that macropinosomes undergo continuous retrograde fusion with subsequent newly-formed macropinosomes and phagosomes challenges the view of maturation as a linear process. Our data indicate a cyclic model whereby nascent macropinosomes fuse with a slightly older population, distinguished by PIPs such as PI(3,5)P_2_. A proportion of these will again fuse with the next round of macropinosomes whilst the rest presumably progress further and lose fusogenicity to enter a terminally digestive phase (**Figure 7**). We show these retrograde fusion events provide a mechanism to rapidly deliver a large volume of hydrolases, V-ATPase and other digestive machinery to newly internalised vesicles.

**Figure 7:**
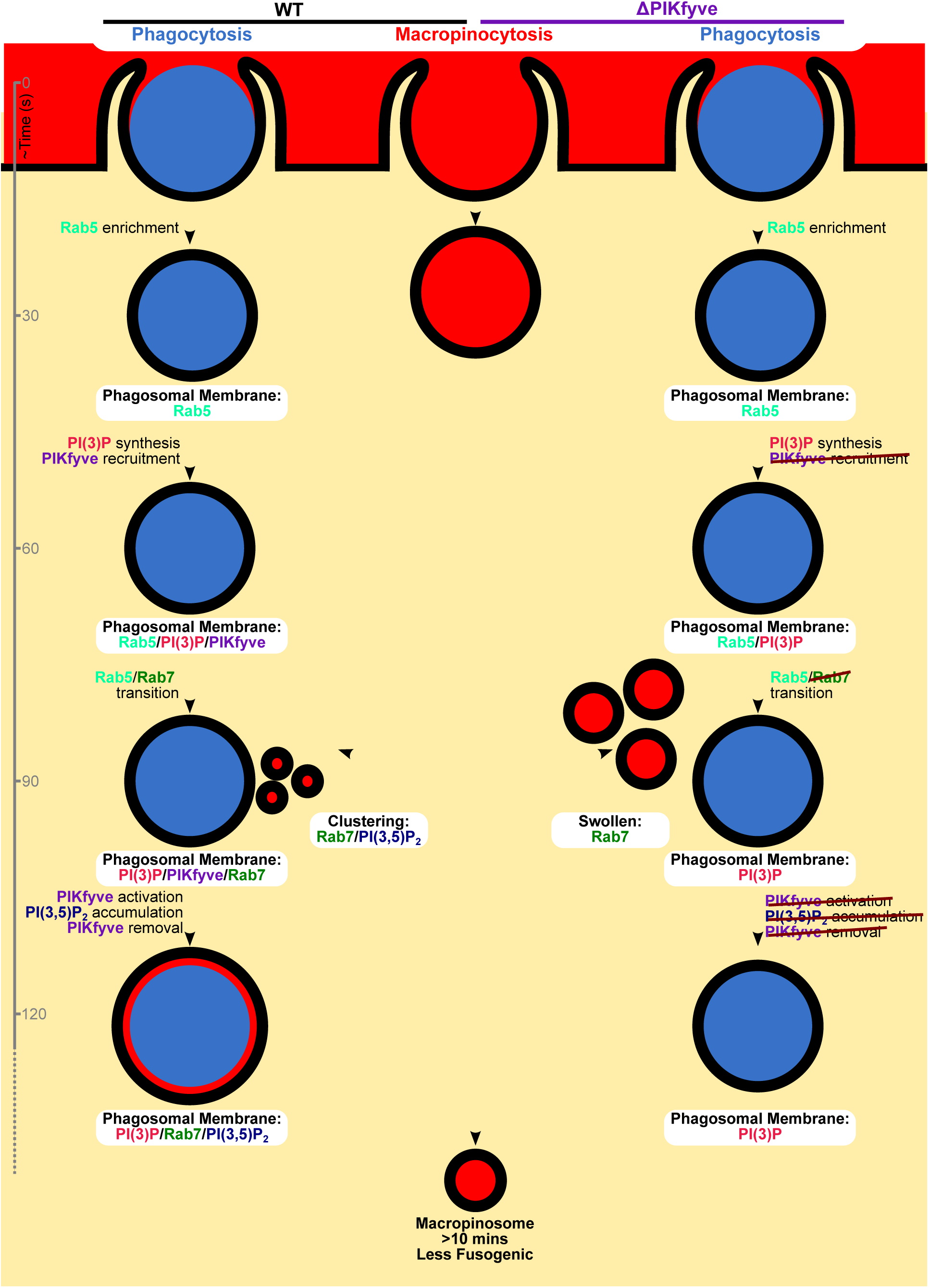
Model for the first 2 minutes of phagosome maturation. The left side indicates the maturation of phagosomes in wild-type cells, with PIKfyve-deficient cells on the right. Macropinosome maturation in the centre. Wild-type phagosomes accumulate each of the proteins and lipids indicated as per Figure 5G, with PIKfyve becoming activated approximately 1 minute after recruitment, at the point that macropinosomes of a similar age fuse and PIKfyve itself dissociates. In the absence of PIKfyve, macropinosomes do not shrink as efficiently, and do not cluster around, or fuse with phagosomes.

Rapid enzyme delivery may be particularly important for both macropinosomes and phagosomes due to their large size and low surface area-to-volume ratio. We find that fusion between macropinosomes and phagosomes depends on PIKfyve, and coincides with the presence of PI(3,5)P_2_ on the macropinosomes. Loss of PIKfyve blocked both the clustering of macropinosomes around phagosomes and the delivery of macropinocytic material. Previous studies have also implicated PIKfyve during endosome fusion events. For example, in mammalian cells fusion of macropinosomes with lysosomes is at least partially regulated by recruitment of the septin cytoskeleton in a PIKfyve-dependent manner (Dolat and Spiliotis, 2016). However, *Dictyostelium* do not possess septins and we find no defect in macropinosomes maturation in these cells, indicating that fusion of phagosomes with lysosomes uses an alternative PIKfyve-dependent mechanism. Nonetheless, PIKfyve appears to have a conserved role in regulating endosomal mixing, and the disruption of fusion between macropinosomes and phagosomes likely accounts for a significant proportion of the acidification and proteolysis defects we previously observed (Buckley et al., 2019).

PIKfyve disruption also results in an almost complete block in Rab7 delivery to phagosomes. In wild-type cells, Rab7A-positive vesicles continually clustered around the surface of nascent phagosomes, with Rab7A intensifying on the phagosomal membrane gradually over several minutes. This indicates that on *Dictyostelium* phagosomes at least, the majority of Rab7 accumulates by fusion with Rab7-positive compartments. This is consistent with recent work (Tu et al., 2022) which also describes fusion of Rab7-positive vesicles with early-Rab5 positive macropinosomes, but contrasts with the canonical view of Rab7 recruitment from a cytosolic pool via the activities of the Mon1/Ccz complex (Langemeyer et al., 2020). Importantly however, although Rab7A is markedly decreased on ΔPIKfyve phagosomes, it is not completely absent with roughly 10% still recruited compared to macropinosomes in the same cell (estimated from our images). Therefore, some Rab7A can accumulate on phagosomes independently of PIKfyve. In the absence of obvious vesicles clustering around the phagosomes, we speculate this small pool of PIKfyve-independent Rab7 comes from a cytosolic pool via classical Rab5-Rab7 exchange. However, this alone is insufficient to for subsequent fusion with other Rab7A-positive compartments, further Rab7 enrichment, and lysosomal fusion.

A strong candidate to mediate the fusion of Rab7 vesicles with phagosomes is the homotypic fusion and protein sorting (HOPS)-tethering complex. In yeast, the Rab7 homolog, Ypt7, directly interacts with the HOPS complex which itself mediates endosomal fusion (Brett et al., 2008; Ostrowicz et al., 2010). A similar, less direct interaction between Rab7 and HOPS occurs in higher eukaryotes through the Rab7 effector RILP (Lin et al., 2014; van der Kant et al., 2013). In *Dictyostelium* disruption of Rab7A also results in severe defects in lysosomal activity and delivery of premature lysosomal enzymes (Rupper et al., 2001). It is possible that this tethering or fusion activity somehow requires PI(3,5)P_2_, but this remains unexplored.

An alternative possibility is through the activity of phagosomal ion channels. PI(3,5)P_2_ has been shown to activate calcium ion channels on phagosomal membranes, and in mammalian cells, over-expression of the PI(3,5)P_2_-activated calcium ion channel TRPML1 partially rescues the characteristic ΔPIKfyve swollen endosome phenotype (Dong et al., 2010). Whether a reduction in swollen endosomes would also restore heterotypic fusion with macropinosomes remains to be tested, although the *Dictyostelium* TRPML1 homolog (MLCN1), does not localize to phagosomes until after post-lysosome transition (Lima et al., 2012), meaning there must be a slightly different mechanism in this organism.

Nearly all clustering PI(3,5)P_2_-positive macropinosomes were also Rab7/PI(3)P-positive, which further supports their identification as macropinosomes less than 8-minutes old and distinct from younger, PI(3,5)P_2_ -negative, compartments. This finding helps to identify multiple pools of Rab7-positive vesicles which fuse with phagosomes and indicates a more complex picture than the canonical view of a single lysosomal population being delivered. Analyses of *Dictyostelium* phagosomes purified at different stages of maturation also showed different proteases were delivered and recovered over time (Gotthardt et al., 2006; Souza et al., 1997). The timescale in our experiments is much shorter, but both studies suggest that multiple endosomal populations are involved in phagosome maturation. The sequential fusion of different endosomal and lysosomal populations was also observed in studies of macropinosome maturation in macrophages (Racoosin and Swanson, 1993) indicating that this is likely a universal phenomenon.

A final interesting aspect of our studies is the clear differences between phagosome and macropinosome maturation in *Dictyostelium*. Though early steps such as the Rab5-Rab7 transition and acquisition of PI(3)P followed by PI(3,5) P_2_ are identical, only phagosomal degradation is strongly affected by loss of PIKfyve. In contrast, PIKfyve-deficient macropinosomes digest normally and still acquire Rab7, even though their shrinkage is reduced - similar to observations in mammalian cells (Freeman et al., 2020; Kerr et al., 2010). PIKfyve therefore appears to play independent roles in shrinkage and degradation, and in *Dictyostelium* at least, only phagosomes require PIKfyve for Rab7 and lysosomal delivery. These differences are likely driven by signalling from phagocytic receptors, but how this is mediated and how it functionally affects killing and digestion remain unclear.

Many aspects of maturation are shared between phagosomes, macropinosomes, and other endocytic pathways, such as classical clathrin-mediated endosomes. These also transition from a Rab5/PI(3)P-positive early form to later compartments demarked by Rab7, V-ATPase and lysosomal components. Degradation of classically endocytosed receptors, autophagosomes and entotic vesicles are also sensitive to PIKfyve inhibition (de Lartigue et al., 2009; Krishna et al., 2016; Qiao et al., 2021). It is therefore likely that at least some of our observations on phagosomes are applicable to other endocytic routes, although the differences with macropinosomes indicate previously unexpected complexity. Nonetheless, in this study we further define the complex sequence of events that occur in the first minutes of a phagosome, and describe key mechanistic insights into how this is regulated by PIKfyve and PI(3,5)P_2_.

## Acknowledgements

The authors thank Huaqing Cai for Rab expression plasmids, Douwe Veltman and Peggy Paschke for their continued development of his expression system and Robert Insall for his support in the early days of this project. JHV was funded by Royal Society grant RGF\EA\180126, JSK is funded by a Royal Society University Research Fellowship URF\R\201036. Imaging work was performed in the Wolfson Light Microscopy Facility at Sheffield.

---

**Figure S1:**
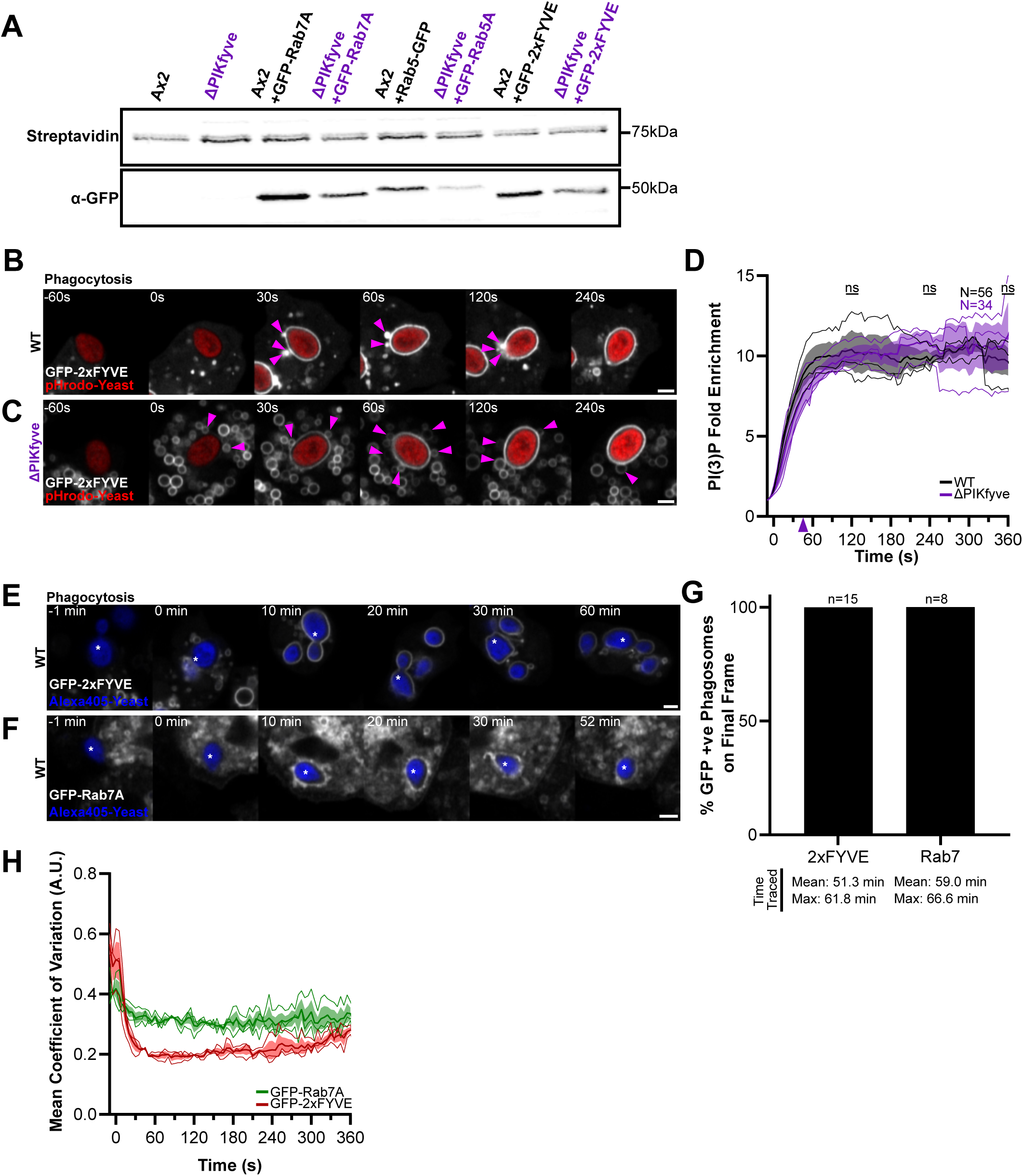
Delivery of reporters to phagosomes and macropinosomes. **(A)** Expression levels of GFP-reporter levels in wild-type and ΔPIKfyve cells. Western blot of whole cell lysates probed with anti-GFP antibody. Streptavidin staining of endogenously biotinylated proteins used as loading control. (**B)** and **(C)** Representative timelapses of the PI(3)P reporter, GFP-2xFYVE, during phagocytosis of pHrodo-yeast (red) in wild-type and ΔPIKfyve cells. Pink arrows indicate GFP-2xFYVE labelled vesicles clustering around the phagosomal membrane from 30s. GFP-2xFYVE enrichment is quantified in **(D)**. There were no statistically significant differences at any timepoints tested. **(E)** and **(F)** Long-term timelapses of GFP-2xFYVE and GFP-Rab7A recruitment to phagosomes. Both reporters remain associated for at least 1 hour after engulfment. The time each event could be tracked varied due to the point in the video when engulfment occurred but the proportion of phagosomes still positive for each reporter at the end of each movie is shown in **(G)**. Only those able to be tracked for >30 minutes were scored. **(H)** Variation in the fluorescent signal around phagosomes of GFP-2xFYVE, and GFP-Rab7A. The coefficient of variation along a linescan around the phagosome was measured at each time point. Graphs show mean coefficients of variation ± SEM. Variation in GFP-Rab7A signal is greater than 2xFYVE, indicating a patchy distribution. All scale bars = 2 μm.

**Figure S2:**
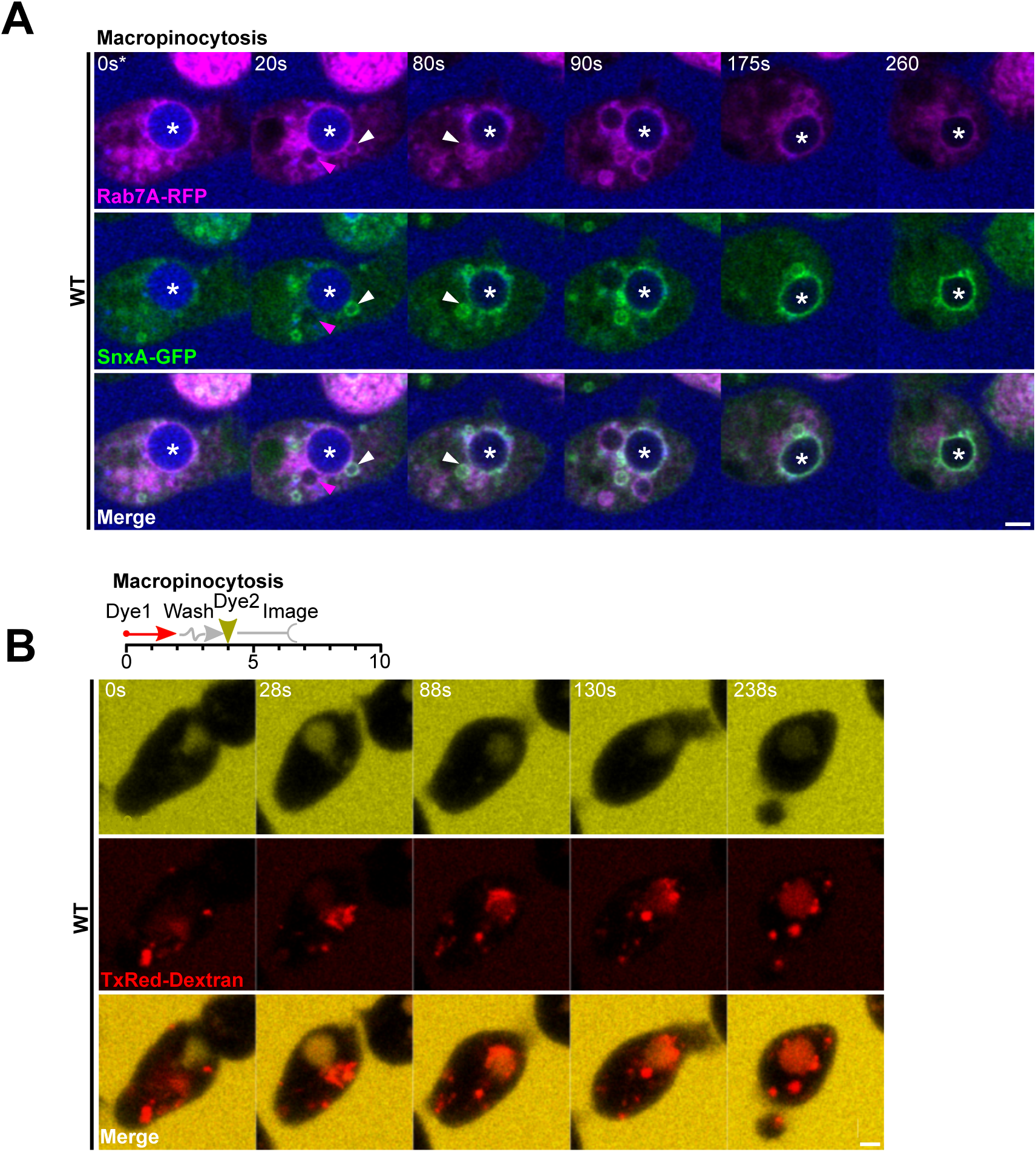
Delivery to Macropinosome. **(A)** Timelapses of RFP-Rab7A (magenta) and SnxA-GFP (green) during macropinocytosis of Alexa405-labelled 70kDa dextran in wild-type cells. Arrowheads indicate clustered SnxA and Rab7 positive vesicles. **(B)** Timelapse of cells after sequential pulses of TxRed and FITC dextran. The older macropinosomes (red) can be seen fusing and mixing contents with a newly formed macropinosome (yellow). All scale bars = 2 μm.

## Video legends

**Video 1:** Simultaneous imaging of RFP-Rab5A and GFP-Rab7A recruitment to phagosomes in wild-type (top) and ΔPIKfyve(bottom) cells. Scale bar = 2 μm

**Video 2:** GFP-Rab5A dynamics at phagosomes in both wild-type (left) and ΔPIKfyve(right) cells. Scale bar = 2 μm

**Video 3:** Enrichment of GFP-Rab5A at phagocytic cups. Multiple failed attempts to complete engulfment of a pHrodo labelled yeast demonstrate that GFP-Rab5A is enriched at the cup before internalisation completes. Scale bar = 2 μm

**Video 4:** PI(3)P dynamics at phagosomes in both wild-type (left) and ΔPIKfyve(right) cell. Cells expressing GFP-2xFYVE reporter for PI(3)P. Scale bar = 2 μm

**Video 5:** GFP-Rab7A dynamics at phagosomes in both wild-type (left) and ΔPIKfyve(right) cells. Scale bar = 2 μm

**Video 6:** PI(3,5) P2 dynamics during phagosome maturation. Wild-type cells expressing the SnxA-GFP probe for PI(3,5) P2. Note small vesicles clustering before membrane enrichment. Scale bar = 2 μm

